# MicroED as a powerful tool for structure determination of macrocyclic drug compounds directly from their powder formulations

**DOI:** 10.1101/2023.07.31.551405

**Authors:** E Danelius, G Bu, H Wieske, T Gonen

**Author notes:** Correspondence to T.G. Denotes equal contribution (E Danelius, G Bu).

## Abstract

Macrocycles are important drug leads with many advantages including the ability to target flat and featureless binding sites as well as act as molecular chameleons and thereby reach intracellular targets. However, due to their complex structures and inherent flexibility, macrocycles are difficult to study structurally and there are limited structural data available. Herein, we use the cryo-EM method MicroED to determine the novel atomic structures of several macrocycles which have previously resisted structural determination. We show that structures of similar complexity can now be obtained rapidly from nanograms of material, and that different conformations of flexible compounds can be derived from the same experiment. These results will have impact on contemporary drug discovery as well as natural product exploration.

## Introduction

The length and complexity of present-day drug discovery has motivated researchers to explore new modalities with more complex structures and target dynamics as compared to conventional rule of 5 therapeutics. [1, 2] Macrocycles, for example, are fascinating drug leads; they interact with their targets in a highly dynamic way, they can be fine-tuned by optimizing the inter-as well as intramolecular interactions, and their flexibility span allows them to act as molecular chameleons and to reach intracellular targets. [3] Macrocycles can be responsive meaning that their conformations can be controlled and switched on and off by using external stimuli. [4] Due to their ability to bind to and modulate ‘undruggable’ targets such as flat protein surfaces or targets with high mutation rate, and with the development of diverse target-tailored libraries of macrocyclic compounds, [5] these ‘beyond rule of 5’ modalities have gained increased interest for biopharmaceutical companies as well as academia. [1, 3] Most macrocycles originate from natural products sources, and as such, their structural determination is key for further optimization. [6] In addition, many oral drug candidates fail in clinical phases due to limitations in physicochemical, mechanical, and pharmacokinetic characterization, where phenomena like structural polymorphism can lead to the complete loss of bioavailability. [7] There are currently 67 FDA approved macrocyclic drugs of which a majority are still lacking atomic resolution structural data. Because of the size and flexibility of macrocycles they are very challenging to crystalize and structurally characterize by X-ray crystallography.

Microcrystal Electron Diffraction (MicroED) [8-10] is a cryogenic electron microscopy (cryo-EM) method capable of determining atomic structures from sub-micrometer sized crystals, as small as a billionth the size as those required for single crystal X-ray diffraction (XRD). [11-13] Due to the many unique advantages of MicroED as compared to other structural methods, including the small amounts of material required, the fact that structures can be obtained rapidly directly from powder formulations, and the possibility of determining structures from compound mixtures, MicroED has been recognized as a new method for contemporary drug discovery. [14-16] In some recent examples MicroED was used to determine the composition of small molecule drug mixtures, [17] the structure of antihistamine levocetirizine which has resisted determination for over 25 years, [18] the novel structure of mirabegron, [19] and new molecular salts of the anti-psychotic drug olanzapine. [20] Although impressive examples, these molecules all fall well within the rule of 5 space with restricted size, flexibility and complexity.

Here we employ MicroED to study large and structurally flexible macrocyclic therapeutics in the beyond rule of 5 space (Figure 1), and thereby addressing the long-standing need for fast and reliable structure determination in natural product and new modalities drug discovery. The smaller and more structurally rigid macrocycles brefeldin A and romidepsin (Figure 1) have been described previously and were used here as proof of concept. The additional macrocycles studied here have resisted structural characterization due to difficulties in growing the large and well-ordered crystals required for XRD. We obtained all structures directly from the commercial powders, i.e. completely bypassing any crystallization assays. The last macrocycle in our series, paritaprevir, is of special interest since it is lacking any accessible structural information such as XRD structure in the Cambridge Crystallographic Data Centre (CCDC), target bound structure in the Protein Data Bank (PDB) or any solution phase NMR data. This clearly shows the expected impact of MicroED in future natural product and beyond rule of five discovery.

**Figure 1.**
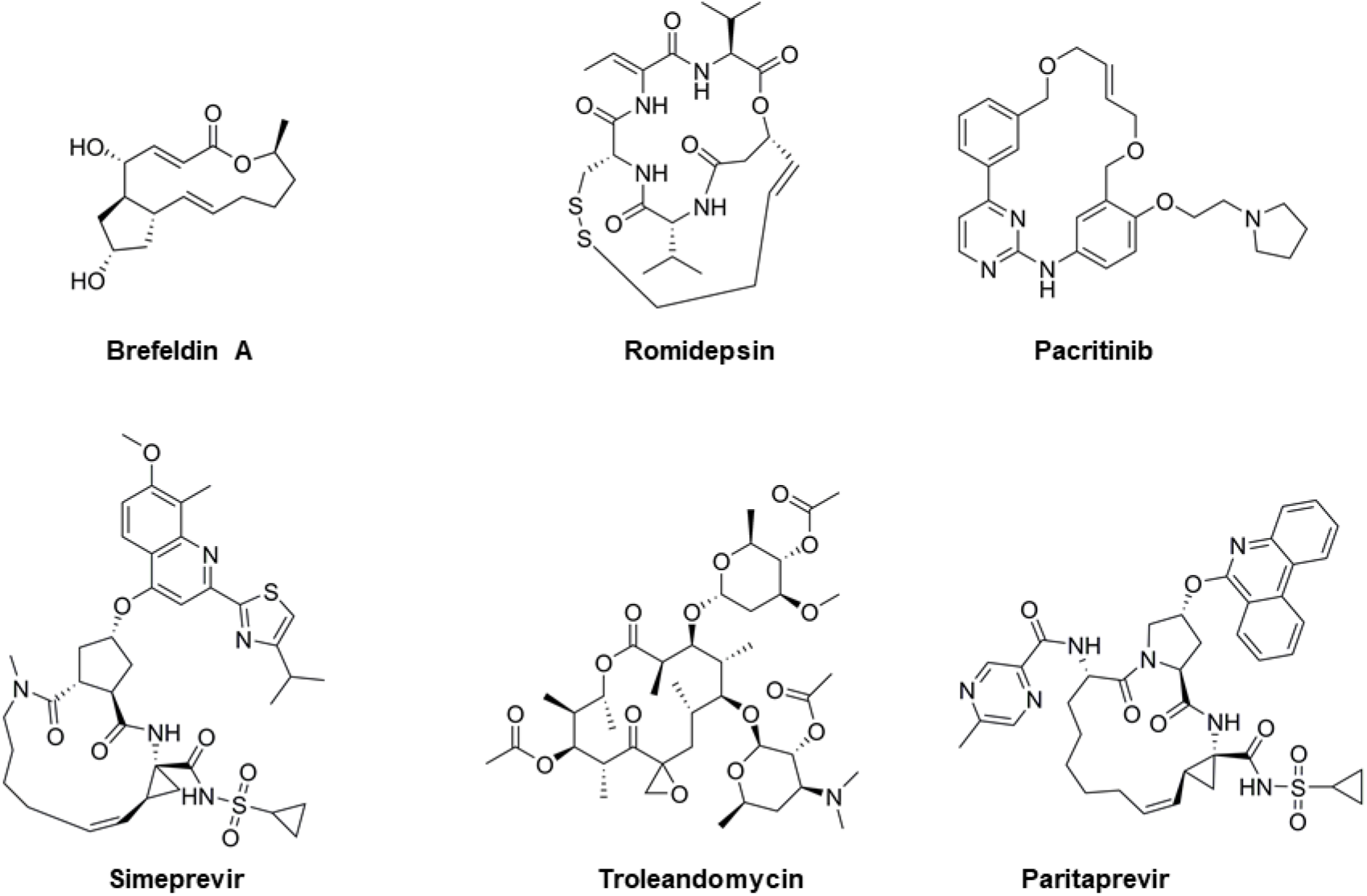
Chemical structures of the macrocyclic drugs investigated by MicroED. A macrocycle is defined as having a cyclic core of 12 heteroatoms or more, giving rise to an increased flexibility in comparison to the more commonly found heterocycles of up to seven atoms.

## Results and discussion

### MicroED grid preparation and diffraction screening

The structural determination of small molecules by MicroED directly from their powders was previously demonstrated. [11, 21, 22] Remarkably, the authors simply applied powder materials to EM grids and collected electron diffraction data directly from the nanocrystalline fragments present in the commercial preparations. For brefeldin A, romidepsin, pacritinib and troleandomycin (Figure 1), a similar grid preparation protocol was applied: the powders were ground between two cover slips and directly added to pre-clipped EM grids, which were frozen and loaded into the transmission electron microscope (TEM). The quality of the grid preparation was evaluated by low magnification TEM images in combination with single exposures in diffraction mode (Figure 2). Brefeldin A, romidepsin, and pacritinib appeared as plate like microcrystals, and troleandomycin as thin needles. The needle microcrystals appeared slightly bent when imaged at a high tilt. For this reason, we switched to using the continuous carbon grids for all our sample preparations, which are more rigid and flat compared to the holy carbon grids typically used for cryo-EM. Simeprevir and paritaprevir have the most complex structures in our series of macrocycles (Figure 1), and different grid preparations had to be screened before sufficient data could be collected. By dissolving small amounts of the simeprevir and paritaprevir powders into minimal amounts of MeOH and letting the solvent evaporate at room temperature for approximately 20 h, thin needle microcrystals with good diffraction could be identified in the TEM (Figure 2).

**Figure 2.**
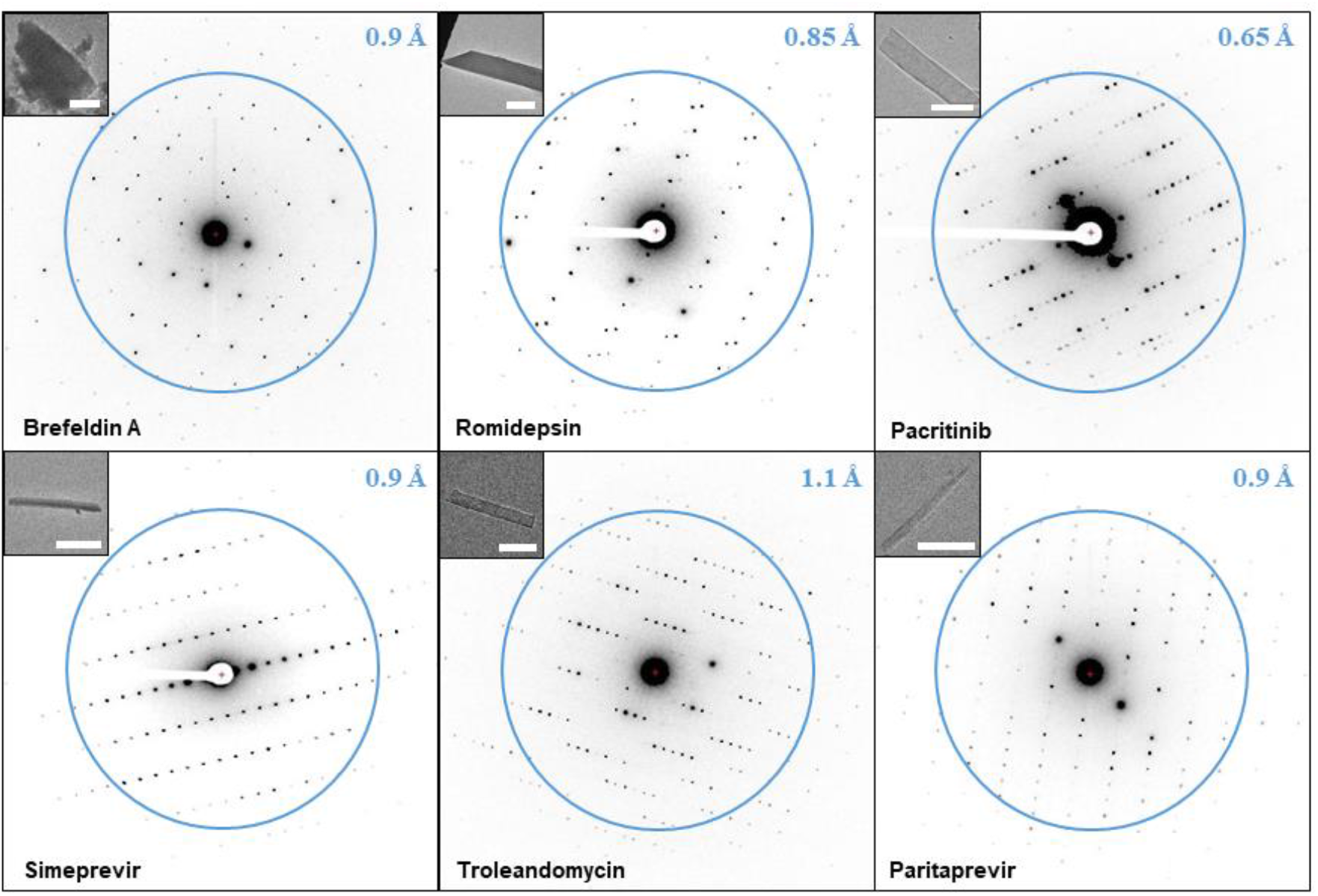
Macrocycle MicroED data. The microcrystal images at 3400x are shown on the top left (the size bar in white corresponds to approximately 5 μm). The resolution is indicated by the blue ring and number.

### MicroED data collection

Initial diffraction screening was performed at 80 K and 200 kV on a scintillator-based Ceta-D detector using a Thermo-Fisher Talos Arctica with EPU-D. The resolution was improved on the Falcon III direct electron detector which has higher detective quantum efficiency (DQE), higher signal-to-noise ratio, and faster readout. [23] This detector was used for all subsequent data collection. The observed diffraction spots from brefeldin A, romidepsin, pacritinib, simeprevir, troleandomycin and paritaprevir extended to 0.8 Å, 0.8 Å, 0.6 Å, 0.9 Å, 1.0 Å and 0.9 Å, respectively (Figure 2). A typical data set was collected as a movie with the sample stage continuously rotating at 0.6 degrees per second, using 2 seconds exposure per frame and an electron dose rate of 0.01 e-/Å^2^/s. By collecting several dataset for each macrocycle, the maximal stage range from -70º to +70º was sampled. With this setup high quality data was collected for pacritinib and simeprevir. For romidepsin and troleandomycin radiation damage was observed less than half minute after initializing data collection, presumably due to the radiation sensitive disulfide bonds [24] and ester groups. [25] Therefore, the exposure was changed to 0.5 seconds per frame as the stage was continuously rotated even faster at 2.0 degrees per second to outrun the damage. Further, for troleandomycin and brefeldin A very few diffraction spots could be observed at the high tilt angles which limited the completeness of the reciprocal space. In addition, due to the preferred orientation on the grid surface adopted by the brefeldin A crystals, generating data with sufficient coverage of the reciprocal space proved challenging even after merging data from multiple crystals. In order to collect a large number of high-quality data sets for merging, a recently developed SerialEM-based high-throughput autonomous MicroED data collection method was employed, [17] where hundreds of MicroED data sets from each sample were automatically generated using the Falcon III detector overnight. Following this protocol more than 200 dataset each were collected for brefeldin A and troleandomycin. Although paritaprevir microcrystals remained stable during data collection and did not show preferred orientation on the grid surface, the majority of the collected data could not be used because of twinning. Hence, SerialEM was used to collect over 800 data sets of paritaprevir of which about 10% showed no twinning. Detailed protocols for initial screening and manual data collection using EPU-D, as well as autonomous data collection using SerialEM, are available in the methods section.

### MicroED data processing and refinement

The continuous-rotation MicroED data were converted to SMV format for data processing using an in-house developed software which is freely available (https://cryoem.ucla.edu/). [26] MicroED data collected using EPU-D was processed using XDS [27] following previously published protocols, [11, 28] and the MicroED data collected using SerialEM was initially processed autonomously including image conversion, indexing, integration and scaling. [17] The *ab initio* structures were solved using either SHELXT [29] or SHELXD [30] in combination with XPREP, followed by refinement in SHELXL. [31] For molecular replacement, data were converted to MTZ format in AIMLESS [32] followed by molecular replacement using Phaser [33], and refinement using Phenix.refine [34]. The pacritinib structure was solved from a single microcrystal at 0.62 Å in Pc (a = 10.52 Å, b = 14.48 Å, c = 15.90 Å, α = γ = 90°, β = 92.944°) using SHELXT, and refined to an R1 value of 0.154 (Table SI-1). The structures of romidepsin and simeprevir were solved by merging 8 manually collected data sets respectively. The romidepsin structure was solved at 0.80 Å with an overall completeness of 81% in P2_1_ (a = 9.17 Å, b = 16.58 Å, c = 9.61 Å, α = γ = 90°, β = 92.952°) using XPREP and SHELXD and refined to an R1 value of 0.229 (Table SI-2). The simeprevir structure was solved at 0.85 Å with an overall completeness of 84% in P1 (a = 5.08 Å, b = 18.69 Å, c = 19.74 Å, α = 89.182°, β = 86.455°, γ = 97.320°) using SHELXT, and refined to a R1 value of 0.132 (Table SI-3). The 0.85 Å dataset for brefeldin A was generated by merging the 4 highest resolution data sets collected by SerialEM, yielding an overall completeness of 93%. The structure was solved in P2_1_2_1_2_1_ (a = 7.51 Å, b = 11.00 Å, c = 19.05 Å, α = β = γ = 90°) using SHELXT, and refined to an R1 value of 0.123 (Table SI-4). For paritaprevir, >600 SerialEM data sets could be processed by the python script, of which 90% showed twinning. The backbone of the macrocycle could be solved from two of the remaining data sets and these two were merged to produce a final dataset with an overall completeness of 89%. The structure was solved at 0.85 Å in P2_1_2_1_2_1_ (a = 5.09 Å, b = 15.61 Å, c = 50.78 Å, α = β = γ = 90°) using XPREP and SHELXD, and refined to an R1 value of 0.147 (Table SI-5). For troleandomycin direct methods failed regardless of whether the data was collected on manually or using SerialEM. Instead, the structure was solved using molecular replacement. The search model was generated using a Monte Carlo Multiple Minimum conformational search followed by molecular mechanics minimization. After multiple trials, a single microcrystal data set processed at 1.70 Å in P2_1_2_1_2_1_ (a = 8.69 Å, b = 23.06 Å, c = 47.33 Å, α = β = γ = 90°), and was refined to R_work_/R_free_ values of 0.27/0.26 (Table SI-6). Detailed protocols for all data processing and structure determination, including crystal and refinement statistics, are available in the methods section.

### Brefeldin A

The antiviral brefeldin A (Figure 1) is a small macrocyclic lactone isolated from the toxic fungi *Penicillium brefeldianum*. It targets the guanine nucleotide exchange factor GBF1, indirectly leading to inhibited protein transport from the endoplasmic reticulum to the golgi complex. [35, 36] Brefeldin A has been evaluated as a lead compound for cancer chemotherapy, however, due to poor solubility, short half-life and significant toxicity it never made it into the clinic. [37] It is one of the most extensively studied macrocycles with four single crystal XRD structures reported in the CCDC database (BREFEL; BREFEL01, BREFEL02; BREFEL03, maximum rmsd=0.0235 Å) and two target bound structures in the pdb database (1RE0 and 1R8Q, rmsd=0.127 Å). As a proof of concept, we determined the structure of brefeldin A directly from the powder formulation using MicroED. Our 0.85 Å structure (Figure 3) compares well to the previously described single crystal structures with the same unit cell dimensions and an average rmsd=0.0434 Å (Figure SI-1a) when comparing macrocyclic heteroatoms. As compared to the target bound structure (Figure SI-1b) the only notable difference is the flipping of the 5 membered ring placing one of the OH-groups in the opposite direction, presumably due to a target interaction of the OH in this position. The crystal packing analysis of brefeldin A reveals each OH to form strong intermolecular hydrogen bonds (Figure SI-1c), leading to a network of hydrogen bonds extending along the crystallographic a axis. Additionally, the covalent bonds between H and O in our MicroED structure are by average 0.251 Å longer than those measured from the structures available in CCDC database that were determined by XRD. This might be due differences between electron and X-ray scattering. [38, 39]

**Figure 3.**
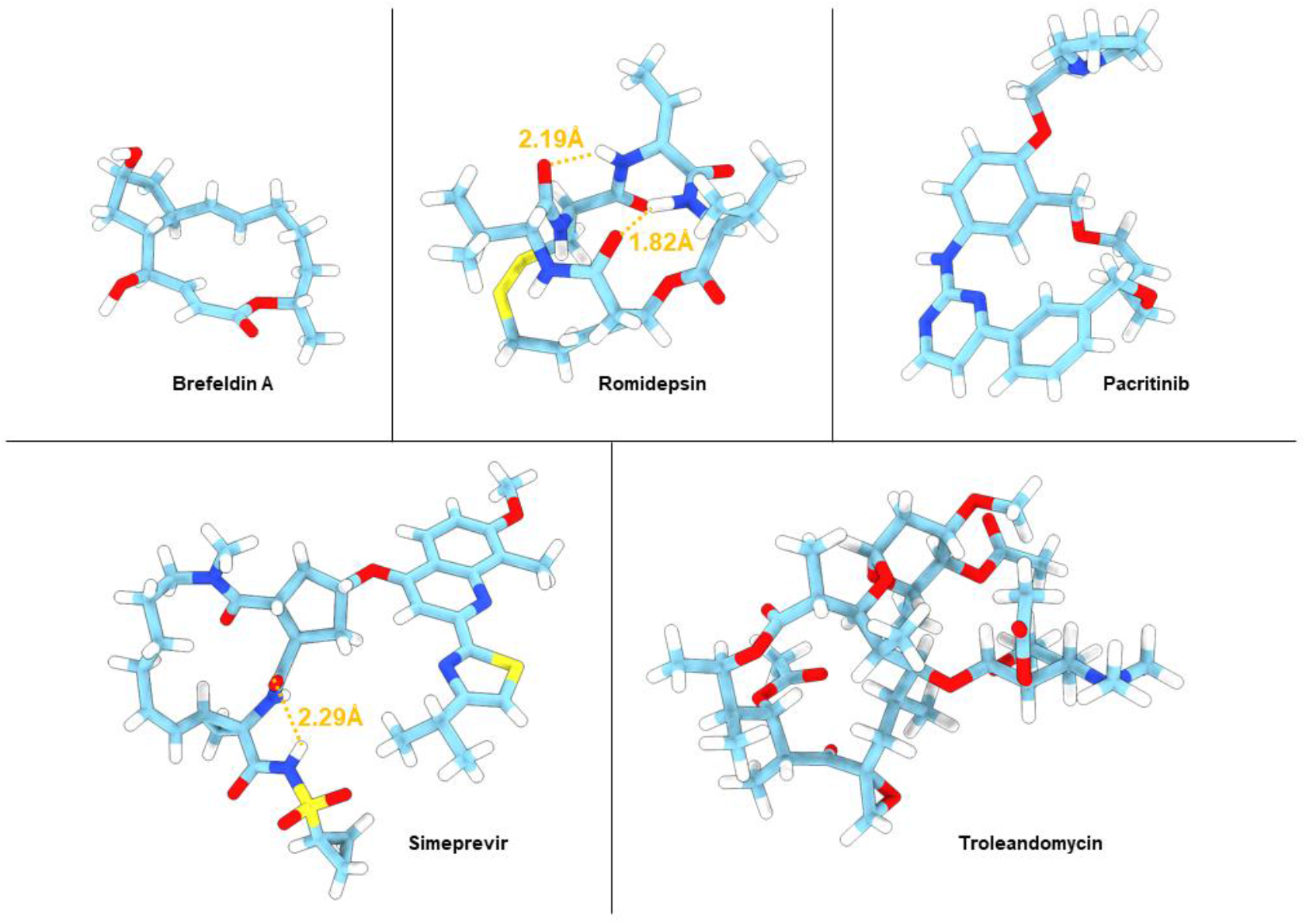
MicroED structures of five of the macrocycles. Atom color: C, light blue; N, dark blue; O, red; H, white. Hydrogen bonds are shown as orange dashed lines.

### Romidepsin

The natural product romidepsin is an anticancer agent isolated from the bacterium *Chromobacterium violaceum*. The structure is composed of a 6-membered cyclic depsipeptide bridged by a 15-membered macrocyclic disulfide ring (Figure 1). In vivo romidepsin acts a pro-drug where cleaving the disulfide bond leads formation of a butenyl thiol, which in turn interacts with zinc in the binding pocket of histone deacetylases leading to cell apoptosis. Romidepsin has been used in clinic for cutaneous T-cell lymphoma since 2009 and other peripheral T-cell lymphomas since 2011. [40] There are limited structural studies of romidepsin with only two entries in the CCDC database (LIDBEF; QEDJOA, rmsd=0.13 Å) and no target bound structures in the pdb database. Our 0.80 Å MicroED structure (Figure 3) is in good agreement with the same unit cell dimension and an average rmsd of 0.1204 Å (Figure SI-2a) when comparing to the macrocyclic heteroatoms. In addition to the intramolecular hydrogen bonds, each molecule forms two intermolecular hydrogen bonds between O and H-N, leading to a network of interactions extending along the crystallographic b axis (Figure SI-2b). Following the initial proof of concept in solving brefeldin A and romidepsin, the scope was shifted towards larger and more flexible macrocycles which had not been previously solved by single crystal XRD.

### Pacritinib

The synthetic macrocyclic anticancer agent Pacritinib (Figure 1), was approved for myelofibrosis in 2022 as the first ever dual inhibitor of Janus kinase 2 (JAK2) and FMS-like receptor tyrosine kinase-3 (FLT3). [41, 42] The only structural data available is that of pacritinib bound to the human quinone reductase 2 (PDB ID: 5LBZ), which is a non-kinase off-target interaction. [43] This structure reveals the three aromatic rings of pacritinib to adopt a fairly flat conformation whereas the carbon chain of the core is folded in the same direction as the sidechain. The 0.62 Å MicroED structure of pacritinib determined here (Figure 3) reveals two conformations in the asymmetric unit, neither of which is similar to the target bound (rmsd= 0.365 Å and 1.41 Å). Interestingly, when comparing the two conformations (rmsd=1.51 Å, Figure SI-3a), the non-aromatic carbon chains of the macrocyclic core are folded in opposite directions with respect to the aromatic rings. Two intermolecular N-H to N hydrogen bonds are identified between the two pyrimidine rings in the asymmetric unit, where one H is located in the difference map. The benzene rings adjacent to hydrogen bonds are tilted in parallel with respect to the hydrogen bonded pyrimidine rings due to their steric crowding effect (Figure SI-3b). The fact that the two conformations determined by MicroED are so different from each other as well as from the quinone reductase bound structure clearly shows that pacritinib is flexible and can adopt various conformations based on the local environment. This chameleonic behavior is important for both the multi-target activity as well as the uptake, and should be optimized for this class of compounds whenever possible based on the structural data.

### Simeprevir

Simeprevir is a serine NS3/4a protease inhibitor used for the treatment of hepatitis C virus (HVC) infections. [44] The only solid-state structural data of simeprevir is the target-bound co-crystal with the serine protease (PDB ID: 3KEE). [45] The binding-site of NS3/4a is rather shallow and flat, making it a difficult to target and at the same time explains the open and flat conformation attained by simeprevir in this structure. Simeprevir has also been studied in solution; NMR ensembles in DMSO and chloroform revealed at least 15 different conformations, where the target-bound crystal structure was not one, again demonstrating the flexibility span and gives an indication as to why it is so difficult to obtain crystals. [46] Similarly as for pacritinib, the 0.85 Å MicroED structure of simeprevir (Figure 3) display two conformations in the asymmetric unit (Figure SI-4a). In this case, however, the conformations are similar (rmsd=0.196 Å) with only some slight difference in side chain orientations. Much like the target-bound crystal structure, the structures adopt open and flat conformations, likely as a result of crystal packing; intermolecular hydrogen bonds form between N-H and O and extend along the crystallographic a axis (Figure SI-4b). Further, the quinoline rings form weak parallel-displaced π-π interactions along the crystallographic a axis. Despite the macrocyclic core being highly similar to the target-bound structures (average rmsd=0.335 Å, Figure SI-1a), the large aromatic moiety adopts a different conformation in the co-crystalized state, and the sulfonamide sidechain displays an extra intramolecular hydrogen bond to the cyclic core (Figure 3).

Simeprevir was developed from a linear peptide precursor, [47] where one of the aims of cyclization is to pre-organize the inhibitor into the bioactive conformation, and thereby lower the entropic penalty required for binding. [48] Since the MicroED structure of simeprevir has the same main core fold as the target bound structure, this might confirm the preorganized state.

### Troleandomycin

The semi-synthetic macrolide troleandomycin is an antibiotic based of the natural product oleandomycin. The structure (Figure 1) consists of a macrocyclic lactone ring with two flexible sugar substituents, one desosamine and one cladinose. Similarly to the macrocycles described above, there is very limited structural data for troleandomycin. In fact, the only published data is that of troleandomycin bond to the ribosomal subunit of *Deinococcus radiodurans* (1OND). [49] From this structure, it was revealed that the troleandomycin binds to the ribosomal subunit near the peptidyl transferase tunnel entrance, in an open and flat conformation with the desosamine and one cladinose in the same plane as the macrocyclic core. As discussed above, the MicroED structure of troleandomycin was obtained using molecular replacement with an input ensemble generated by a Monte Carlo Multiple Minimum conformational search. From our calculated ensemble troleandomycin was observed to adopt open conformations where the desosamine and one cladinose are oriented in the same plans as the core, open conformations where the sugars are on opposite sides of the core, and closed conformations with the sugars on top of each other, similar to a sandwich structure. For the MicroED structure, the sugars adopts a planar conformation similar to the target bound structure, however, macrocyclic core adopts a slightly different fold (Figure SI 1-5).

### Paritaprevir

Paritaprevir is a first-generation inhibitor of the HVC NS3/4a protease, and one of the 25 highest molecular weight drugs approved for oral administration. [50, 51] It has been studied by calculations and powder diffraction, where it was found to have a substantial conformational flexibility, but no crystal structures are accessible for comparison. [52] The flexibility of paritaprevir is essential both for the uptake and for binding to the relatively flat and featureless HCV protease binding site, but similarly to the other macrocycles discussed here this leads to challenges in obtaining the structural data. In addition, paritaprevir has a high number of hydrogen bond donors and acceptors, and low chemical stability due to oxidative liability. Our 0.85 Å structure displays a relatively open conformation with an intramolecular hydrogen bond between the secondary sulfonamide group and the macrocyclic core (Figure 4a). The 3D-PSA of the MicroED structure as computed by Schrodinger QikProp (174 Å^2^) compares well to the speculated open conformation (201 Å^2^), as well as the predicted target bound conformation (186 Å^2^). [52] The packing analysis of the paritaprevir structure reveals intermolecular hydrogen bonds between the carbonyl O and secondary NH on the macrocyclic core, extending along the crystallographic a axis (Figure SI-6). Further, solvent accessible channels form along the crystallographic a axis (Figure 4 b and c). From the macrocycles described herein, the observation of water channels is unique to paritaprevir. It is known that flexible molecules with chameleonic behavior can display erratic aqueous solubility and form crystal structures with large voids that can accommodate a significant amount of water, which can be crucial for their solubility, adsorption and bioavailability.

**Figure 4.**
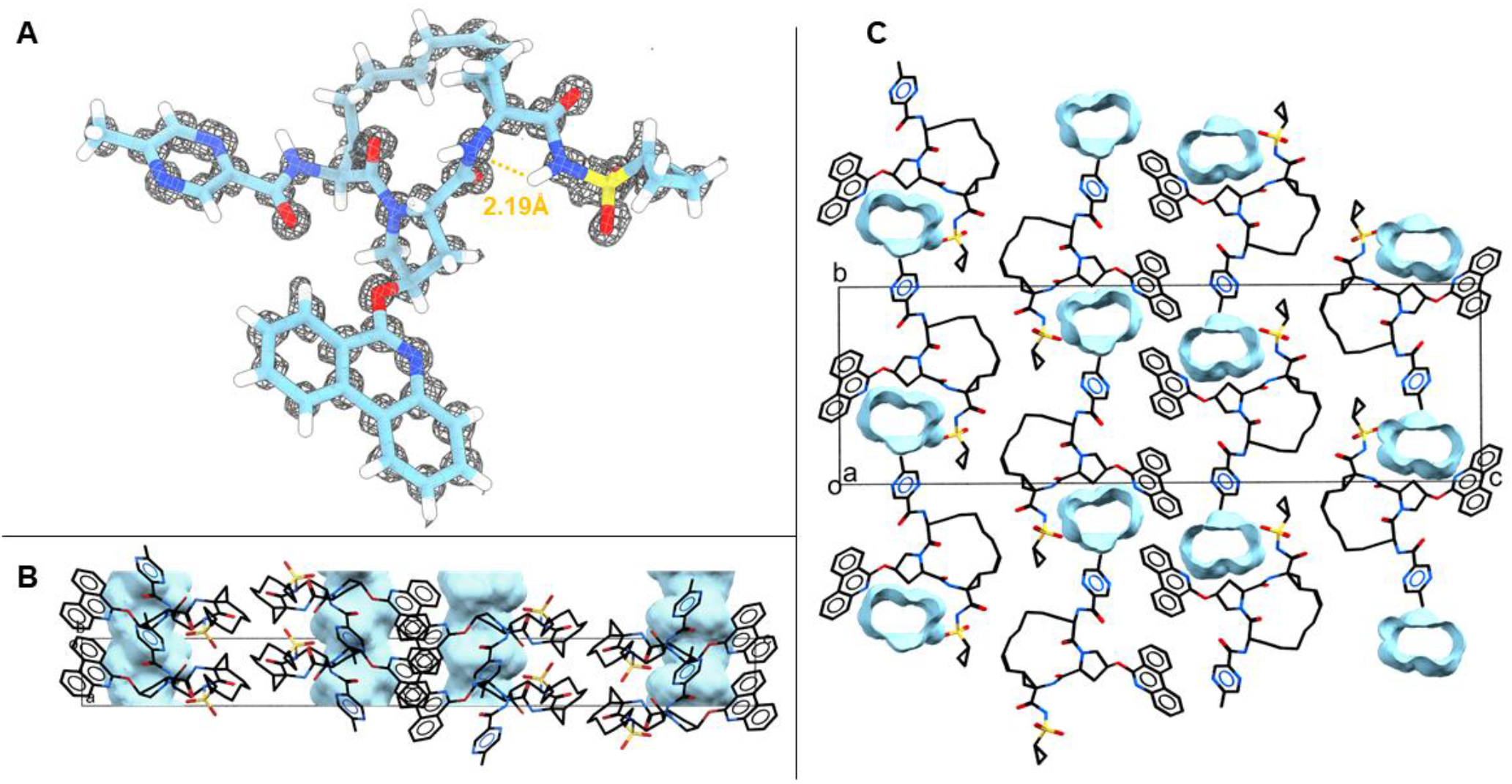
Structure and packing of paritaprevir. **A**. MicroED structure with the 2Fo-Fc map in grey mesh. Atom color: C, light blue; N, dark blue; O, red; H, white. Hydrogen bonds are shown as orange dashed lines. **B**. Side view of crystal packing. **C**. Top view of crystal packing. Unit cell (box) of solved crystal structure in P2_1_2_1_2_1_ space group packing. Atom color: C, black; N, blue; O, red. Hydrogen atoms are omitted for clarity. Solvent/water channels indicated in blue in B and C.

## Conclusion

The importance of optimization in the beyond Ro5 space constituting macrocyclic and flexible molecules for oral drugs was recently reemphasized. [53, 54]. By the use of MicroED we have determined the atomic structures of several macrocycles directly from nanograms of powder material, without any prior crystallization, thereby giving access to previously unattainable structures. Despite their wide use, many of the macrocycles described herein were previously not described in the unbound state due to difficulties in growing the appropriate size crystals for XDR. Further, by the use of MicroED different conformations could be identified from the same experiments. From the novel structure of paritaprevir, we could identify large solvent channels, which can impact the probability for crystallization as well as the drug formulation. Either being provided by nature or invented *de novo*, the optimization of potent, cell-permeable, and orally available macrocyclic drugs has many unknowns to be resolved. This is especially true for the design of macrocyclic molecular chameleons which is still poorly understood. We show that MicroED can considerably impact future research in this field.

## Acknowledgements

This study was funded in part by the National Institutes of Health P41GM136508. Portions of this research or manuscript completion were developed with funding from the Department of Defense grants MCDC-2202-002. Effort sponsored by the U.S. Government under Other Transaction number W15QKN-16-9-1002 between the MCDC, and the Government. The US Government is authorized to reproduce and distribute reprints for Governmental purposes, notwithstanding any copyright notation thereon. The views and conclusions contained herein are those of the authors and should not be interpreted as necessarily representing the official policies or endorsements, either expressed or implied, of the U.S. Government. The PAH shall flowdown these requirements to its subawardees, at all tiers. The Gonen laboratory is supported by funds from the Howard Hughes Medical Institute. E.D. thanks The Wenner-Gren Foundations for their support through the Wenner-Gren Postdoctoral Fellowship. L.W. thanks The Bengt Lundqvist Memorial Foundation, the Liljewalch foundation, and the Bergmarks foundation for fellowship support.

## Methods and materials

All the macrocycle compounds are commercially available. Brefeldin A was purchased from MedChemExpress, romidepsin and troleandomycin from Focus Biomolecules, simeprevir and paritaprevir from Invivochem, and pacritinib from Sigma Aldrich.

### Sample preparation

For brefeldin A, romidepsin, pacritinib, and troleandomycin, approximately 0.5 mg of each compound was crushed between two cover slips, transferred to glass vials, and applied to pre-clipped continuous carbon 400-mesh copper TEM grids (Ted Pella Inc.) by gently shaking the compounds together with the grids in the vials. For simeprevir and paritaprevir, approximately 1 mg of each compound was dissolved into methanol. The vials were left open in a fume hood at room temperature for approximately 20 h. The resulting deposits were scraped from the glass walls and applied to pre-clipped TEM grids as described above. All grids were prepared by negative glow-discharging the for 30 seconds on each side at 15 mA in a PELCO easiGlow (Ted Pella Inc.) prior to mixing with compounds. The grids were pre-frozen in liquid nitrogen before loading them into the microscope. **MicroED screening and manual data collection**. Microcrystals on the TEM grids were screened using EPU-D (Thermo Fisher Scientific) on a Thermo-Fisher Talos Arctica electron microscope operating at 80 K with an acceleration voltage of 200 kV, corresponding to a wavelength of 0.0251 Å. The whole grid atlases were acquired in the “Atlas” settings at a magnification of 210x. Microcrystals were imaged in the “Search/Eucentric height” settings at a magnification of 3,400x. Upon identification of single microcrystals, the selected area aperture (approximately 2 µm in diameter) was inserted to cover the target area, and the microscope was switched to the “Diffraction Acquisition” settings for taking still diffraction images under the parallel electron beam conditions (C2 lens intensity of 45.2% inserted with an aperture size of 70). Once sharp and high-resolution diffraction spots were observed, manual MicroED data was collected as movies on the Falcon III detector as the stage was continuously rotating. For pacritinib and simeprevir, the data was recorded at a rate of 2 second exposure per frame and a tilt speed of 0.6 degree per second. For romidepsin, the data was recorded at a rate of 0.5 second exposure per frame and a tilt speed of 2 degree per second. **High-throughput automated MicroED data collection using SerialEM**. High-throughput MicroED data were automatically collected on Falcon III detector using SerialEM. [17] The whole grid atlas was acquired as low-magnification montage at a magnification of 155x. Grid squares containing microcrystals were selected by the SerialEM “Navigator”, and acquired for medium-magnification montages at a magnification of 2,600x. During medium-magnification montage, the “Fine eucentricity” function in SerialEM was selected to assign the eucentric heights for each grid square in the corresponding maps. Microcrystals were picked from each medium-magnification montage within the “Navigator” window for data collection. MicroED data collection was performed in the SerialEM “Record” mode where the microscope was set for the parallel electron diffraction settings (C2 lens intensity of 45.2% inserted with an aperture size of 20, resulting in the beam size of approximately 1.5 µm in diameter). An in-house developed script was used for the automatic MicroED data collection within the set tilt ranges. For troleandomycin, the script was set to collect data at an acquisition rate of 0.5 second exposure per frame and a stage tilt speed of 2 degree per second.

For brefeldin A and paritaprevir, the script was set to collect data at an acquisition rate of 1 second exposure per frame and a stage tilt speed of 1 degree per second.

### Data processing and structure determination

Romidepsin, pacritinib, and simeprevir data collected in the manual mode were converted to images in SMV format using an in-house developed software which is freely available (https://cryoem.ucla.edu/microed). Images were processed in XDS [27] for indexing, integration, and scaling. A single crystal dataset of pacritinib was converted to SHELX format in XDS. For romidepsin and simeprevir, 8 datasets each were merged and converted to SHELX format in XDS, respectively. Brefeldin A, troleandomycin, and paritaprevir data from the SerialEM collection were processed using the in-house developed python script, [17] and the high-quality datasets identified were manually reprocessed in XDS. For structure determination, 4 brefeldin A datasets, and 2 paritaprevir datasets were merged and converted to SHELX formats in XDS, respectively. A single crystal dataset of troleandomycin was converted to MTZ format in AIMLESS. [32] The *ab initio* structures of brefeldin A, pacritinib, and simeprevir were solved by SHELXT [7] followed by structure refinement in SHELXL. [31] The reflection files of romidepsin and paritaprevir were prepared by XPREP (Bruker) and their *ab initio* structures were determined by SHELXD, [30] followed by structure refinement in SHELXL. The troleandomycin structure was phased by molecular replacement using Phaser [33] and refined using phenix.refine. [34]

## Notes

### Competing Interest Statement

The authors have declared no competing interest.

