## Supplemental for "MicroED as a powerful tool for structure determination of macrocyclic drug compounds directly from their powder formulations"


Danelius E[1,2*]., Bu G[2*], Wieske H[3] and Gonen T[1,2,4$]

1. Howard Hughes Medical Institute, University of California Los Angeles, Los Angeles, CA 90095, USA.
2. Department of Biological Chemistry, University of California Los Angeles, 615 Charles E.Young Drive South, Los Angeles, CA 90095, USA.
3. Department of Chemistry – BMC, Uppsala University, Husargatan 3, 75237 Uppsala, Sweden.
4. Department of Physiology, University of California Los Angeles, 615 Charles E. Young Drive South, Los Angeles, CA 90095, USA.

* Denotes equal contribution


**Supplementary Data Table 1**. MicroED data collection and refinement statistics for pacritinib.

| | |
|---|---|
| Stoichiometric formula | $C_{28}H_{32}N_4O_3$ |
| Radiation wavelength (Å) | 0.0251 |
| Temperature (K) | 80 |
| Number of crystals | 1 |
| Crystal description | plate |
| Resolution (Å) | 0.62 |
| Crystal system | monoclinic |
| Space group | Pc |
| a (Å) | 10.52 |
| b (Å) | 14.48 |
| c (Å) | 15.90 |
| α (°) | 90 |
| β (°) | 92.944 |
| ɣ (°) | 90 |
| Z | 4 |
| Total reflections | 28,160 |
| Unique reflections | 9,857 |
| $R_{obs}$ (%) | 9.1 |
| $R_{meas}$ (%) | 11.2 |
| I/σI | 6.17 |
| $CC_{1/2}$ (%) | 99.5 |
| Completeness (%) | 89.9 |
| $R_1$ | 0.1537 |
| $wR_2$ | 0.4132 |
| GooF | 1.162 |
| CCDC number | |

**Supplementary Data Table 2**. MicroED data collection and refinement statistics for romidepsin.

| | |
|---|---|
| Stoichiometric formula | $C_{24}H_{36}N_4O_6S_2$ |
| Radiation wavelength (Å) | 0.0251 |
| Temperature (K) | 80 |
| Number of crystals | 8 |
| Crystal description | plate |
| Resolution (Å) | 0.80 |
| Crystal system | monoclinic |
| Space group | $P2_1$ |
| a (Å) | 9.17 |
| b (Å) | 16.58 |
| c (Å) | 9.61 |
| α (°) | 90 |
| β (°) | 92.952 |
| ɣ (°) | 90 |
| Z | 2 |
| Total reflections | 11,059 |
| Unique reflections | 2,513 |
| $R_{obs}$ (%) | 23.8 |
| $R_{meas}$ (%) | 26.6 |
| I/σI | 4.97 |
| $CC_{1/2}$ (%) | 98.1 |
| Completeness (%) | 81.0 |
| $R_1$ | 0.2293 |
| $wR_2$ | 0.4488 |
| GooF | 1.248 |
| CCDC number | |

**Supplementary Data Table 3.** MicroED data collection and refinement statistics for simeprevir.

| | |
|---|---|
| Stoichiometric formula | $C_{38}H_{47}N_5O_7S_2$ |
| Radiation wavelength (Å) | 0.0251 |
| Temperature (K) | 80 |
| Number of crystals | 8 |
| Crystal description | needle |
| Resolution (Å) | 0.85 |
| Crystal system | triclinic |
| Space group | P1 |
| a (Å) | 5.08 |
| b (Å) | 18.69 |
| c (Å) | 19.74 |
| α (°) | 89.182 |
| β (°) | 86.455 |
| ɣ (°) | 97.320 |
| Z | 2 |
| Total reflections | 22,621 |
| Unique reflections | 5,294 |
| $R_{obs}$ (%) | 16.0 |
| $R_{meas}$ (%) | 18.2 |
| I/σI | 5.43 |
| $CC_{1/2}$ (%) | 98.4 |
| Completeness (%) | 84.2 |
| $R_1$ | 0.1324 |
| $wR_2$ | 0.3424 |
| GooF | 1.030 |
| CCDC number | |

**Supplementary Data Table 4**. MicroED data collection and refinement statistics for brefeldin A.

| | |
|---|---|
| Stoichiometric formula | $C_{16}H_{24}O_4$ |
| Radiation wavelength (Å) | 0.0251 |
| Temperature (K) | 80 |
| Number of crystals | 4 |
| Crystal description | plate |
| Resolution (Å) | 0.85 |
| Crystal system | orthorhombic |
| Space group | $P2_12_12_1$ |
| a (Å) | 7.51 |
| b (Å) | 11.00 |
| c (Å) | 19.05 |
| α (°) | 90 |
| β (°) | 90 |
| ɣ (°) | 90 |
| Z | 4 |
| Total reflections | 10,217 |
| Unique reflections | 1,471 |
| $R_{obs}$ (%) | 17.2 |
| $R_{meas}$ (%) | 18.6 |
| I/σI | 7.68 |
| $CC_{1/2}$ (%) | 98.8 |
| Completeness (%) | 92.9 |
| $R_1$ | 0.1229 |
| $wR_2$ | 0.3368 |
| GooF | 1.200 |
| CCDC number | |

**Supplementary Data Table 5**. MicroED data collection and refinement statistics for paritaprevir.

| | |
|---|---|
| Stoichiometric formula | $C_{40}H_{43}N_7O_7S$ |
| Radiation wavelength (Å) | 0.0251 |
| Temperature (K) | 80 |
| Number of crystals | 2 |
| Crystal description | needle |
| Resolution (Å) | 0.85 |
| Crystal system | orthorhombic |
| Space group | $P2_12_12_1$ |
| a (Å) | 5.09 |
| b (Å) | 15.61 |
| c (Å) | 50.78 |
| α (°) | 90 |
| β (°) | 90 |
| ɣ (°) | 90 |
| Z | 4 |
| Total reflections | 15,086 |
| Unique reflections | 3,607 |
| $R_{obs}$ (%) | 20.2 |
| $R_{meas}$ (%) | 23.2 |
| I/σI | 5.20 |
| $CC_{1/2}$ (%) | 98.9 |
| Completeness (%) | 89.0 |
| $R_1$ | 0.1472 |
| $wR_2$ | 0.4080 |
| GooF | 1.156 |
| CCDC number | |

**Supplementary Data Table 6**. MicroED data collection and refinement statistics for troleandomycin.

| | |
|---|---|
| Stoichiometric formula | $C_{41}H_{67}NO_{15}$ |
| Radiation wavelength (Å) | 0.0251 |
| Temperature (K) | 80 |
| Number of crystals | 1 |
| Crystal description | needle |
| Resolution (Å) | 1.70 |
| Crystal system | orthorhombic |
| Space group | $P2_12_12_1$ |
| a (Å) | 8.69 |
| b (Å) | 23.06 |
| c (Å) | 47.33 |
| α (°) | 90 |
| β (°) | 90 |
| ɣ (°) | 90 |
| Z | 4 |
| Total reflections | 3,162 |
| Unique reflections | 1,011 |
| $R_{obs}$ (%) | 12.1 |
| $R_{meas}$ (%) | 16.5 |
| I/σI | 6.3 |
| $CC_{1/2}$ (%) | 98.3 |
| Completeness (%) | 85.5 |
| $R_{work}$ | 0.26 |
| $R_{free}$ | 0.27 |

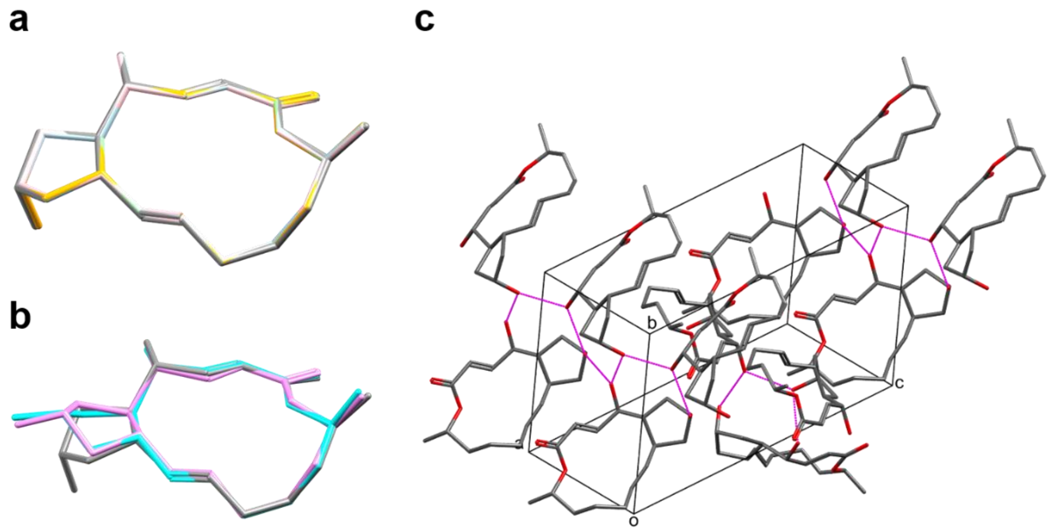


**Figure SI-1. Structure and packing of brefeldin A. a.** Overlay of the heavy atoms of the macrocyclic rings of the MicroED structure (gray) with the SCXRD structures: CCDC codes BREFEL[1] (pink, rmsd=0.0509 Å), BREFEL02[2] (orange, rmsd=0.0384 Å), BREFEL03[3] (light green, rmsd=0.0394 Å), and BREFEL04[4] (light blue, rmsd=0.0448 Å). **b.** Overlay of the heavy atoms of the macrocyclic rings of the MicroED structure (gray) with the target bound structures: PDB codes 1RE0[5] (cyan, rmsd=0.106 Å), and 1R8Q[6] (violet, rmsd=0.114 Å). **c.** Crystal packing of brefeldin A with the unit cell box shown in black. Atom colors: C, gray; O, red. Intermolecular hydrogen bonds (1.68 Å and 1.75 Å) are shown in magenta. Hydrogen atoms are omitted for clarity.

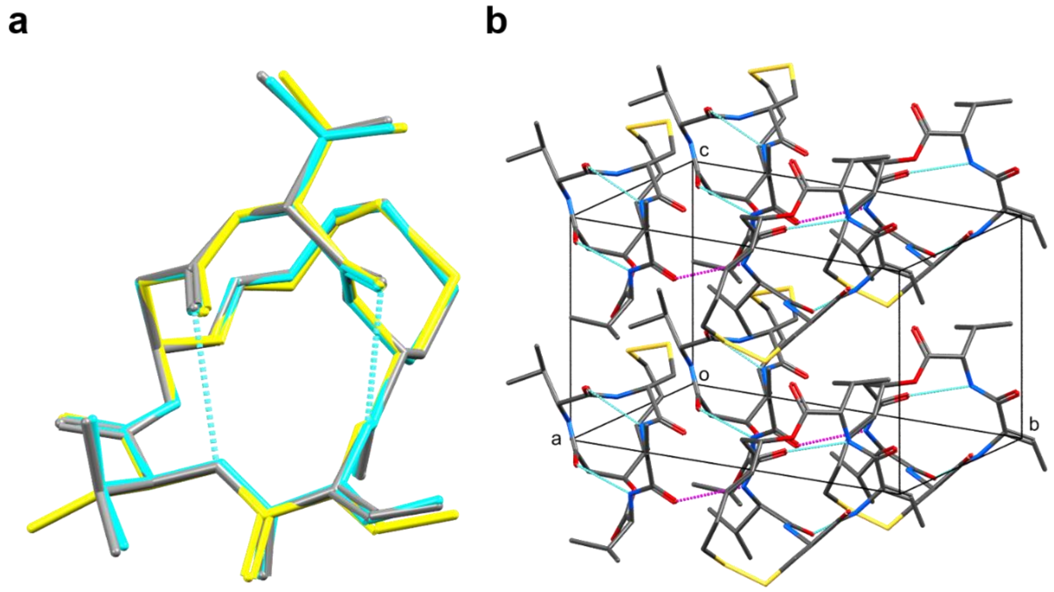


**Figure SI-2. Structure and packing of romidepsin. a.** Overlay of the heavy atoms of the macrocyclic rings of the MicroED structure (gray) with the SCXRD structures: CCDC codes QEDJOA[7] (cyan, rmsd=0.0977 Å) and LIBDEF[8] (yellow, rmsd=0.143 Å). Intramolecular hydrogen bonds (2.16 Å and 2.36 Å) are shown in cyan dashed lines. **b.** Crystal packing of romidepsin with the unit cell box is shown in black. Atom colors: C, gray; N, blue; O, red; S, yellow. Intramolecular hydrogen bonds (2.16 Å and 2.36 Å) are shown in cyan dashed lines. Intermolecular hydrogen bonds (2.21 Å) are shown in magenta. Hydrogen atoms are omitted for clarity.

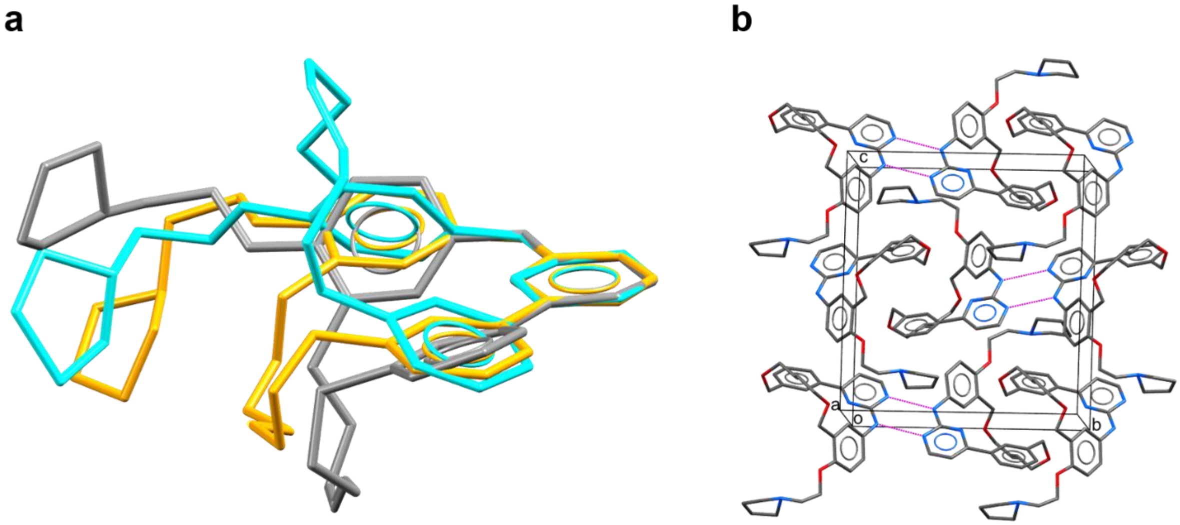


**Figure SI-3. Structure and packing of pacritinib. a.** Overlay of the biphenyl aromatic atoms of the macrocyclic rings of the two MicroED conformations (conformer 1, gray; conformer 2, cyan; rmsd=1.51 Å when comparing macrocyclic heteroatoms) and the target bound structure (PDB ID 5LBZ[9] orange, rmsd=0.365 Å and 1.41 Å compared with conformer 1 and 2, respectively). **b.** Crystal packing of pacritinib with the unit cell box is shown in black. Atom colors: C, gray; N, blue; O, red. Intermolecular hydrogen bonds (1.90 Å and 2.10 Å) are shown in magenta. Hydrogen atoms are omitted for clarity.

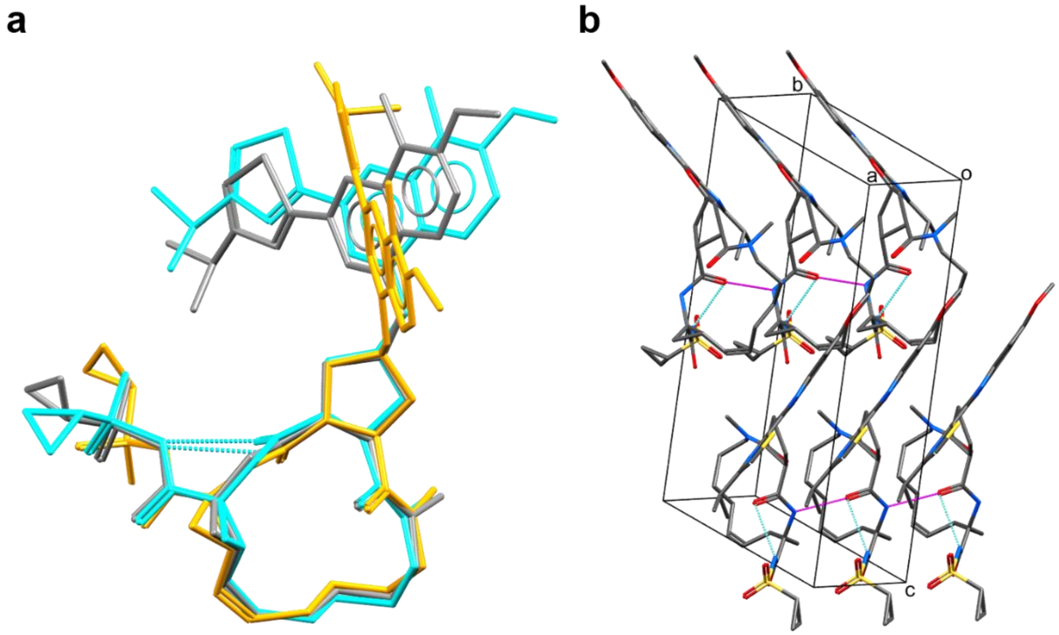


**Figure SI-4. Structure and packing of simeprevir. a.** Overlay of the heavy atoms of the macrocyclic rings of the two MicroED conformations (conformer 1, gray; conformer 2, cyan; rmsd=0.196 Å) and the target bound structure (PDB ID: 3KEE[10], orange, rmsd=0.264 Å and 0.406 Å compared with conformer 1 and 2, respectively). Intramolecular hydrogen bonds (2.05 Å conformer 1, and 2.29 Å conformer 2) shown in cyan dashed lines. **b.** Crystal packing of simeprevir with the unit cell box is shown in black. Atom colors: C, gray; N, blue; O, red; S, yellow. Intramolecular hydrogen bonds (2.05 Å conformer 1, and 2.29 Å conformer 2) are shown in cyan dashed lines. Intermolecular hydrogen bonds (2.29 Å conformer 1, and 2.24 Å conformer 2) are shown in magenta. Hydrogen atoms are omitted for clarity.

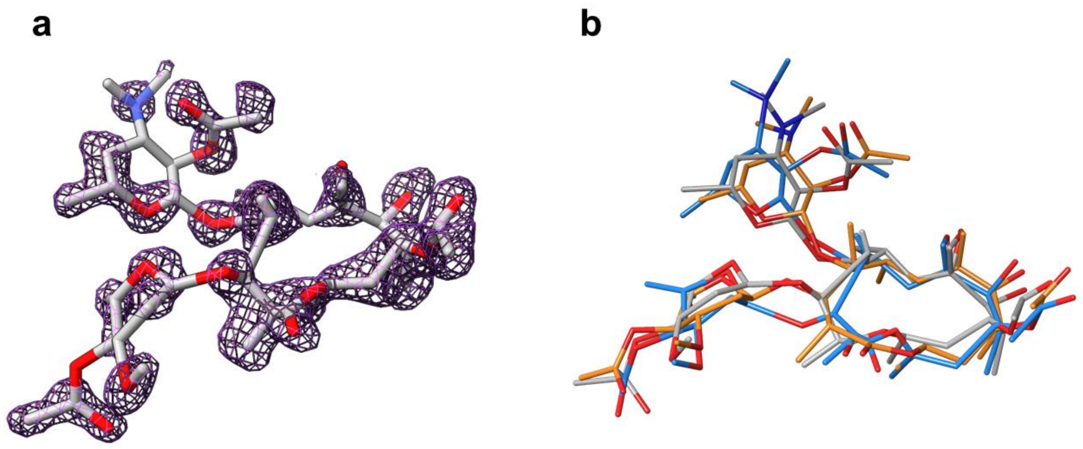


**Figure SI-5. Structure of troleandomycin. a.** MicroED structure with the 2Fo-Fc map in grey mesh (2σ). Two components were found in the molecular replacement, conformer 1 shown here. Atom colors: C, gray; N, blue; O, red. Hydrogen atoms are omitted for clarity. **b.** Overlay of the heavy atoms of the macrocyclic rings of the two MicroED conformations (conformer 1, gray; conformer 2, blue) and the target bound structure (PDB ID: 1OND[11], orange).

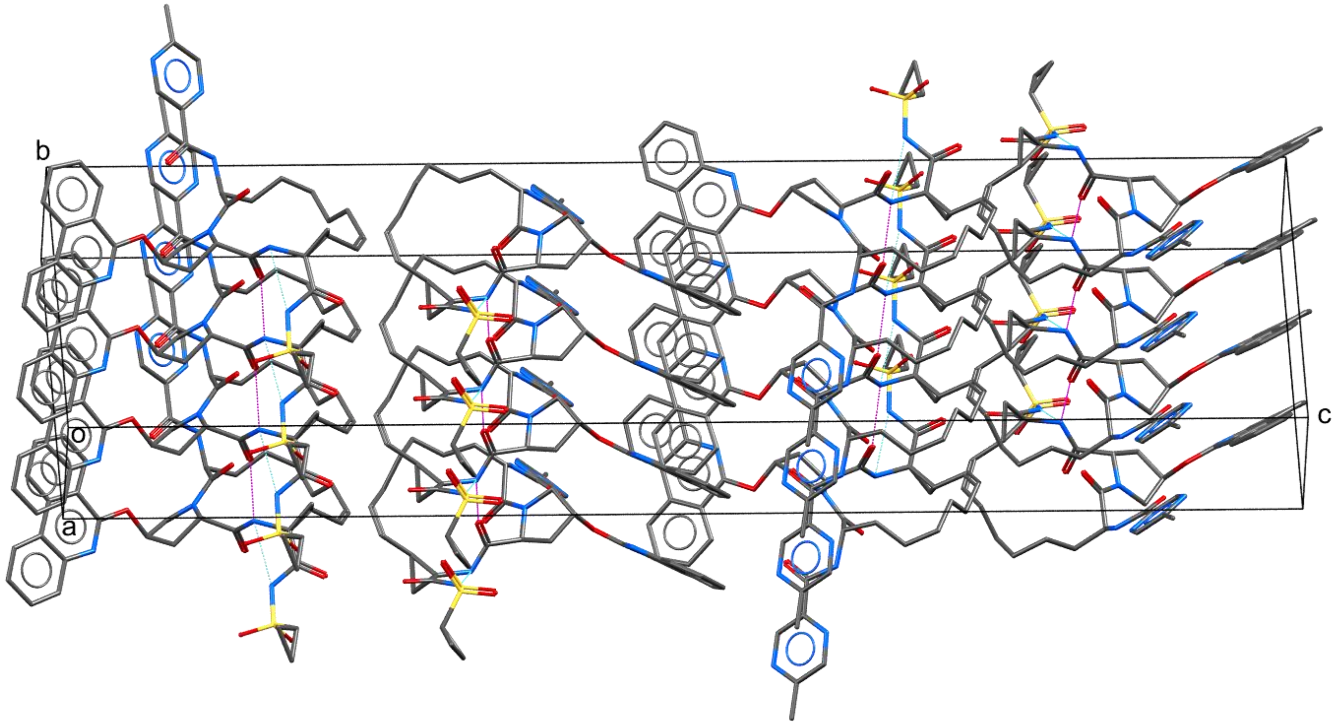


**Figure SI-6. Packing of paritaprevir.** Crystal packing of paritaprevir with the unit cell box is shown in black. Atom colors: C, gray; N, blue; O, red; S, yellow. Intramolecular hydrogen bonds (2.19 Å) are shown in cyan dashed lines, and intermolecular hydrogen bonds (2.26 Å) are shown in magenta. Hydrogen atoms are omitted for clarity.
